# Single nuclei chromatin profiling of ventral midbrain reveals cell identity transcription factors and cell type-specific gene regulatory variation

**DOI:** 10.1101/2020.06.10.144626

**Authors:** Yujuan Gui, Kamil Grzyb, Mélanie H. Thomas, Jochen Ohnmacht, Pierre Garcia, Manuel Buttini, Alexander Skupin, Thomas Sauter, Lasse Sinkkonen

## Abstract

Cell types in ventral midbrain are involved in diseases with variable genetic susceptibility such as Parkinson’s disease and schizophrenia. Many genetic variants affect regulatory regions and alter gene expression. We report 20 658 single nuclei chromatin accessibility profiles of ventral midbrain from two genetically and phenotypically distinct mouse strains. We distinguish ten cell types based on chromatin profiles and analysis of accessible regions controlling cell identity genes highlights cell type-specific key transcription factors. Regulatory variation segregating the mouse strains manifests more on transcriptome than chromatin level. However, cell type-level data reveals changes not captured at tissue level. To discover the scope and cell-type specificity of *cis*-acting variation in midbrain gene expression, we identify putative regulatory variants and show them to be enriched at differentially expressed loci. Finally, we find TCF7L2 to mediate *trans*-acting variation selectively in midbrain neurons. Our dataset provides an extensive resource to study gene regulation in mesencephalon.

## INTRODUCTION

The ventral midbrain, or mesencephalon, is one of the most evolutionary conserved brain structures in mammals (1). It is involved in tasks such as processing of sensory information and eliciting motor and cognitive control through dopaminergic circuits (1). It is of particular interest due to its involvement in human diseases like Parkinson’s disease and schizophrenia, whose development and progression are significantly influenced by individual’s genetic susceptibility (2–5).

Like other brain regions, midbrain harbors many different cell types that exhibit both functional and molecular diversity (6–8). A cell type can be distinguished by its gene expression profile. Transcriptomic analysis at single cell level has identified 20 cell types and 58 subtypes in ventral midbrain (8). These unique gene expression profiles defining cell state and cellular identity are controlled by epigenetic mechanisms and achieved by dynamic interplay between chromatin and expressed transcription factors (TFs). In particular, regulation by cell type-specific master regulators, TFs that open and specifically bind to gene regulatory regions, results in distinct gene expression profiles between cell types (9). The chromatin landscape and accessibility of TF binding sites can be elucidated using epigenomic analysis such as the assay for transposase-accessible chromatin followed by sequencing (ATAC-seq) (10). So far, the ability to isolate pure populations of various brain cell types has been limiting the progress in the field. However, recent developments in single nuclei chromatin assays have now enabled massive parallel analysis of cell type-specific chromatin profiles in their native context (11–14).

Typical human genomes differ from each other on average by 5 million genetic variants (15). Vast majority of these are located in the non-coding genome and those associated with complex traits are enriched at accessible gene regulatory regions in a cell type-specific manner (16). Genetic variation at regulatory regions can influence TF binding and thereby gene expression either in *cis* or in *trans*, hereafter referred to as gene regulatory variation (17). Identifying genes, regulatory regions and cell types affected by regulatory variants can help to understand the molecular mechanisms underlying the trait in question. C57BL/6J and A/J are two genetically distinct inbred mouse strains often used in neurobiology and to study complex genetic traits. The two strains segregate by ~6 million variants, comparable to the genetic variation between typical human individuals, making them an interesting model system to understand the effects of regulatory genetic variation on the phenotypic expression of complex traits. Indeed, the two strains show genetic differences also in traits associated with midbrain function. For example, A/J is more anxious and less social (18) and has lower motor activity (19). We have recently shown that the two strains exhibit significant differences in their ventral midbrain transcriptomes (20), but the underlying gene regulatory changes, as well as the cell type-specific epigenomic profiles of mouse ventral midbrain, are not known.

Here we performed chromatin accessibility profiling of mouse ventral midbrains from C57BL/6J and A/J at single nuclei level (snATAC-seq). We identify >260 000 individual regulatory regions across 20 658 epigenomic profiles which can distinguish ten main cell types in ventral midbrain, and define sets of unique cell identity genes and identify TFs controlling their expression. Comparison of gene expression and chromatin accessibility between the mouse strains shows that genetically driven differences are more striking at the transcriptomic than chromatin accessibility level. Nevertheless, regulatory regions with alterated chromatin accessibility are enriched at differentially expressed genes and can reveal cell type-specific gene regulation. We find *cis*-acting variants to be enriched at differentially expressed genes and pinpoint the extent of cell type-specific gene regulatory variation. Finally, we suggest canonical Wnt signalling to be a mediator of *trans*-acting variation in midbrain neurons.

## MATERIALS AND METHODS

### Animals

All experimental procedures in this study were in compliance with the European Communities Council Directive 2010/63/EU, following the 3 Rs’ requirements for Animal Welfare. We used two mouse strains in this study, C57BL/6J and A/J, purchased respectively from Charles River and Jackson Laboratory. The study cohorts were bred in-house at the Animal Facility of University of Luxembourg (Esch-sur-Alzette, Luxembourg) and the protocol was approved by the Animal Experimentation Ethics Committee (AEEC) according to the national guidelines of the animal welfare law in Luxembourg (*Règlement grand-ducal* adopted on January 11^th^, 2013). All mice were housed with a 12-hour light–dark schedule and had free access to food and water.

For each mouse, intracardiac perfusion with PBS was performed after anesthesia with a ketamine-medetomidine mix (150 and 1 mg/kg, respectively). The brain was extracted and both midbrains of each mouse were dissected, immediately snap-frozen, stored at −80°C, and used for single cell partitioning as described below.

### Nuclei isolation

For bulk ATAC-seq, frozen midbrains were minced in a Dounce Tissue Grinder (Sigma, D8939-1SET) with A pestle ~10 times following B pestle ~20 times in lysis buffer [5 mM CaCl_2_ (Merck, A546282), 3 mM Mg(Ac)_2_ (Roth, P026.1), 10 mM Tris pH 7.8, 0.17 mM β-mercaptoethanol (Gibco, 21985-023), 160 mM sucrose (Sigma, S0389), 0.05 mM EDTA, 0.05% NP40 (Sigma, I3021)]. The lysate was layered on 3 mL sucrose cushion [1.8 M sucrose, 3 mM Mg(Ac)2, 10 mM Tris pH 8.0, 0.167 mM β-mercaptoethanol], following ultra-centrifugation 30,000g for 1 h (Rotor:Beckman Coulter, MLS-50) at 4°C. After centrifugation, the supernatant was discarded and the nuclei pellet was suspended in resuspension buffer (with 0.1% Tween-20, 0.01% digitonin and 0.1% NP40).

For snATAC-seq, the isolation of nuclei was done according to 10X Genomics protocol CG000212 (Rev A) with minor modification. In brief, frozen midbrain sections were minced in a Dounce Tissue Grinder (Sigma, D8939-1SET) with A pestle ~10 times following B pestle ~20 times in 500 uL chilled lysis buffer [10mM Tris-HCL (pH7.4), 10mM NaCl, 3mM MgCl_2_, 0.1% Tween-20, 0.1% NP40, 0.01% Digitonin, 1% BSA]. The homogenized lysate was incubated on ice for 5 min, following pipette mixing 10x and incubated again on ice for 10 min. Chilled Wash Buffer (500 uL) [10mM Tris-HCl (pH 7.4), 10mM Nacl, 3mM MgCl_2_, 1% BSA, 0.1% Tween-20] was added to the lysed cells and pipetted mix 5x. The lysate was layered on 3 mL sucrose cushion [1.8 M sucrose, 3 mM Mg(Ac)2, 10 mM Tris pH 8.0, 0.167 mM β-mercaptoethanol], following ultra-centrifugation 30,000xg for 1 h (Rotor:Beckman Coulter, MLS-50) at 4°C. After centrifugation, the supernatant was discarded and the nuclei pellet was suspended in Nuclei Buffer (10x PN: 2000153) provided by 10X Genomics Chromium Single Cell ATAC Reagent Kit. The nuclei suspension was passed through a 40 um Flowmi Cell Strainer (Sigma, BAH136800040-50EA).

### Bulk ATAC-seq

Tagmentation of mouse midbrain samples was done based on the OMNI-ATAC supplementary protocol 1 (21) with minor changes. Briefly, 25,000 nuclei were resuspended in 50 μL resuspension buffer (with 0.1% Tween-20, 0.01% digitonin and 0.1% NP40) and lysed for 3 minutes on ice. After washing in 1 mL resuspension buffer (1% Tween-20) samples were centrifuged for 10 minutes at 500xg at 4 °C and supernatant carefully removed. Pellets were resuspended in 25 μL tagmentation mix (Tagment DNA buffer from Illumina, #15027866) containing 2.5 μL Tagment DNA Enzyme (Illumina #15027865) and incubated for 45 min at 37 °C and 1000 rpm in Eppendorf ThermoMixer. Tagmented chromatin was isolated using Zymo Research DNA Clean & Concentrator kit (ZymoResearch ZY-D4013) and eluted in 21 μL elution buffer. Library pre-amplification was done for 5 cycles using primers Ad1 and Ad2.16 (10). Five additional cycles of library amplification were done as determined by qPCR (21). Library cleanup was done using Zymo Research DNA Clean & Concentrator kit followed by AMPure XP bead (Beckman Coulter #A63880) size selection. A first bead incubation using 0.55x volume of beads removes large fragments. After separation of beads on magnetic stand, supernatant was transferred to a fresh tube and incubated for 5 minutes in 1.5x volumes of beads. After washing with 80% Ethanol, beads were resuspended in 20 μL elution buffer. After separation on magnetic stand eluate was transferred to a fresh tube. Library quality was assessed using Agilent DNA High sensitivity Bioanalyzer chip (Agilent #5067-4626).

### Bulk ATAC-seq data analysis

The sequencing of ATAC-seq libraries was done at the sequencing platform in the Luxembourg Centre for Systems Biomedicine (LCSB) of the University of Luxembourg. The paired-end, unstranded library sequencing was performed using Illumina NextSeq 500/550 75 cycles High Output Kit. Raw FASTQ files and BAM files were processed as described in the Materials and Methods section for ChIP-seq. After processing of the BAM files, the peaks were called by Genrich (https://github.com/jsh58/Genrich) with parameters “-r -m 30 -j” to remove PCR duplicates and include only reads with mapping quality of at least 30. Footprints were called by HINT-ATAC (22) with default parameters. Raw FASTQ files were deposited in ArrayExpress with the accession number E-MTAB-8333.

### Single nuclei ATAC-seq library preparation and sequencing

The single nuclei ATAC-seq was performed according to Chrominum Single Cell ATAC Reagent Kits User Guide (CG000168 RevB) with Chromium Single Cell E Chip Kit (10X, 100086), Chromium Single Cell ATAC Library & Gel Bead Kit (10X, 1000111), Chromium Single Cell ATAC Gel Bead Kit (10X, 1000085), Chromium Single Cell ATAC Library Kit (10X 100087), Chromium i7 Multiplex Kit N Set (1000084), Dynabeads MyOne Silane (2000048). In brief, nuclei suspension was loaded with a targeted recovery rate of 10 000 nuclei per sample. snATAC-seq libraries quality were assessed using Agilent DNA High sensitivity Bioanalyzer chip (Agilent #5067-4626) and further sequenced on a 150 cycles High Output Kit using Illumina NextSeq™ 500 with targeted sequencing depth of 25 000 read pairs per nucleus. Raw FASTQ files were deposited in ArrayExpress with the accession number E-MTAB-9225.

### Single nuclei ATAC-seq analysis pipeline

#### Cell ranger

The alignment and filtering were done according to 10X running pipelines. In brief, the fastq files was generated from Illumina sequencer’s base call files, which were later used as inputs to align (MAPQ > 30), filter barcode multiplets and generate accessibility counts for each cell in a single library. The technical replicates for each mouse strain were aggregated to create a single peak-barcode matrix. Each unique fragment is associated with a single cell barcode.

#### Clustering

The snATAC-seq downstream analysis was performed by Signac (version: 0.2.4) in R. The gene activity matrix was calculated with reads in gene body and 2 kb upstream of TSS as a proxy for gene expression. Nuclei with counts less than 5000 were filtered out. The dimensionality was calculated with latent semantic index (LSI) on peaks with at least 100 reads across all cells, which was used as input to generate UMAP graphs.

#### Cluster annotation

The annotation of snATAC-seq clusters took use of the existing scRNA-seq data on midbrains from adult 3-month-old C57BL/6N mice. The anchors were found between the gene activity matrix of snATAC-seq and the top 5000 variable features of scRNA-seq. The cell labels were transferred from the scRNA-seq to the snATAC-seq with normalization on anchor weights calculated from the LSI dimensional reduction, resulting in 10 347 nuclei of C57BL/6J and 10368 nuclei of A/J annotated.

### scRNA-seq data analysis pipeline

#### Data processing

The DGE (digital gene expression) and cell annotation for midbrains of 3-month-old C57BL/6N mice were downloaded from DropViz. The data analysis was performed by Seurat (version: 3.1.4) in R. Only cells with feature counts between 400 to 7000 and being single or well-curated were used in downstream analysis, resulting in 19 967 cells in total. The DGE was natural-log transformed and normalized to mitochondrial read counts. The dimentional reduction was done with UMAP. The clusters were annotated with existing annotation from DropViz.

#### Selection of cell type-identity genes

100 cells were randomly selected from each cell cluster. DGE from each cell type was constructed according to corresponding barcodes. The 85th percentile expression for each gene was calculated on the selected cells. The criteria for cell type-identity genes was defined as: For a particular gene, 60% of cells in a cell type have expression larger than the gene specific 85^th^ percentile; while at most 40% of cells in all other cell types have expression larger than the 85^th^ percentile. This process were repeated for 100 times. Genes that appeared more than 30 times out of 100 were defined as the cell type-identity genes.

#### Defining diffeiential peaks

Differential peaks were defined for each cell type using Wilcoxon rank-sum test (implemented in R package “presto” 1.0.0). The binary peak count matrix from scATAC-seq was normalized by the number of peaks presented in each cell. Then the differential peak analysis was performed with presto on the normalized peak count matrix. Peaks with FDR < 0.05 and log fold-change (logFC) in top 1 % are defined as differential peaks.

### Motif enrichment analysis

#### Geneiating cell type specific bam files

The cell type specific bamfiles were generated by samtools. The barcodes from each replicate of a strain in bamfile was relabled to avoid barcode collapse. After relabelling, the bamfiles from replicates of a sample were merged. The bamfile for each cell type were subtracted based on corresponding barcodes.

#### Peak calling

The peak calling was done by MACS2 (2.1.2) (23) with custom cutoff on p-values according to cutoff analysis with parameters ‘macs2 callpeak --cutoff-analysis’. The ideal cutoff was chosen based on that the selected p-value would not lead to exponential increase of peak numbers.

#### Motif enrichment analysis

The motif enrichment analysis was performed by HOMER (4.11.1) (24) with parameters ‘findMotifsGenome.pl -size given -mask’.

### Chromatin Immunoprecipitation (ChIP)

ChIP was performed on the dissected snap frozen mouse ventral midbrain tissue as previously described (20). Each reaction had 10 – 14 μg of chromatin and 10% aliquot was used as input DNA. Immunoprecipitation was performed overnight with 5 μL of H3K27ac antibody (Millipore, 17-614) at 4°C with rotation.

### ChIP-seq data analysis

The sequencing of the chromatin samples was done at the sequencing platform in the LCSB of the University of Luxembourg. The single-end, unstranded sequencing with read length of 75 bp was performed with Illumina NextSeq 500 machine. FastQC (v0.11.5) was used for raw reads quality assessment (25). Generation of BAM files, including steps of adapter removal, mapping and duplicate marking, were done with PALEOMIX pipeline (v1.2.12) (26), followed by mapping with BWA (v.0.7.16a) (27). Backtrack algorithm applied the quality offset of Phred score to 33. Duplicate reads were marked and the mouse reference genome, GRCm38.p5 (mm10, patch 5) was downloaded from GENCODE (https://www.gencodegenes.org/). Finally, validation of the Bam files was done using Picard (v2.10.9) (28). Raw FASTQ files were deposited in ArrayExpress with the accession number E-MTAB-8333.

The H3K27ac (H3 lysine 27 acetylation) ChIP-seq peaks, enhancers and super-enhancers, were called by HOMER (24) with default parameters. The signal normalization in pairwise comparison was done by THOR (v0.10.2) (29), with TMM normalization and adjusted p-value cut-off 0.01.

## RESULTS

### Single nuclei chromatin profiles of ventral midbrain and identification of major cell types in two mouse strains

To unravel the cell type-specific gene regulation in midbrain, and how it impacts genetic regulatory variation, we performed ATAC sequencing at single nuclei level (snATAC-seq) on dissected midbrains from two genetically distinct mouse strains, C57BL/6J and A/J (Figure 1, Supplementary Figure S1). Perfused ventral midbrain sections from both strains were used for the partitioning and barcoding with a total of 13 640 and 13 259 nuclei from C57BL/6J and A/J, respectively, and following high throughput sequencing. After filtering of multiplets and nuclei with low coverage, approximately 290 million reads per mouse strain were retained, corresponding to 10 298 (C57BL/6J) and 10 360 (A/J) individual accessibility profiles (Figure 1A). The bulk chromatin accessibility profile aggregated across single nuclei (bulk snATAC-seq) from C57BL/6J showed a total of 231 390 peaks. Notably, 99.7% of regular bulk ATAC-seq peaks obtained from an independent C57BL/6J midbrain section overlapped with bulk snATAC-seq peaks (Figure S1). Moreover, the bulk snATAC-seq profile from A/J with 235 157 peaks was also highly correlated with C57BL/6J profile (Pearson R > 0.97). Finally, to distinguish accessible regions at enhancers and promoters actively engaged in transcriptional control, we performed ventral midbrain bulk level ChIP-seq analysis in both mouse strains for histone H3 lysine 27 acetylation (H3K27ac) (30, 31). Both bulk ATAC-seq and H3K27ac ChIP-seq showed clear correlation with midbrain gene expression levels (Figure S2).

**Figure 1.**
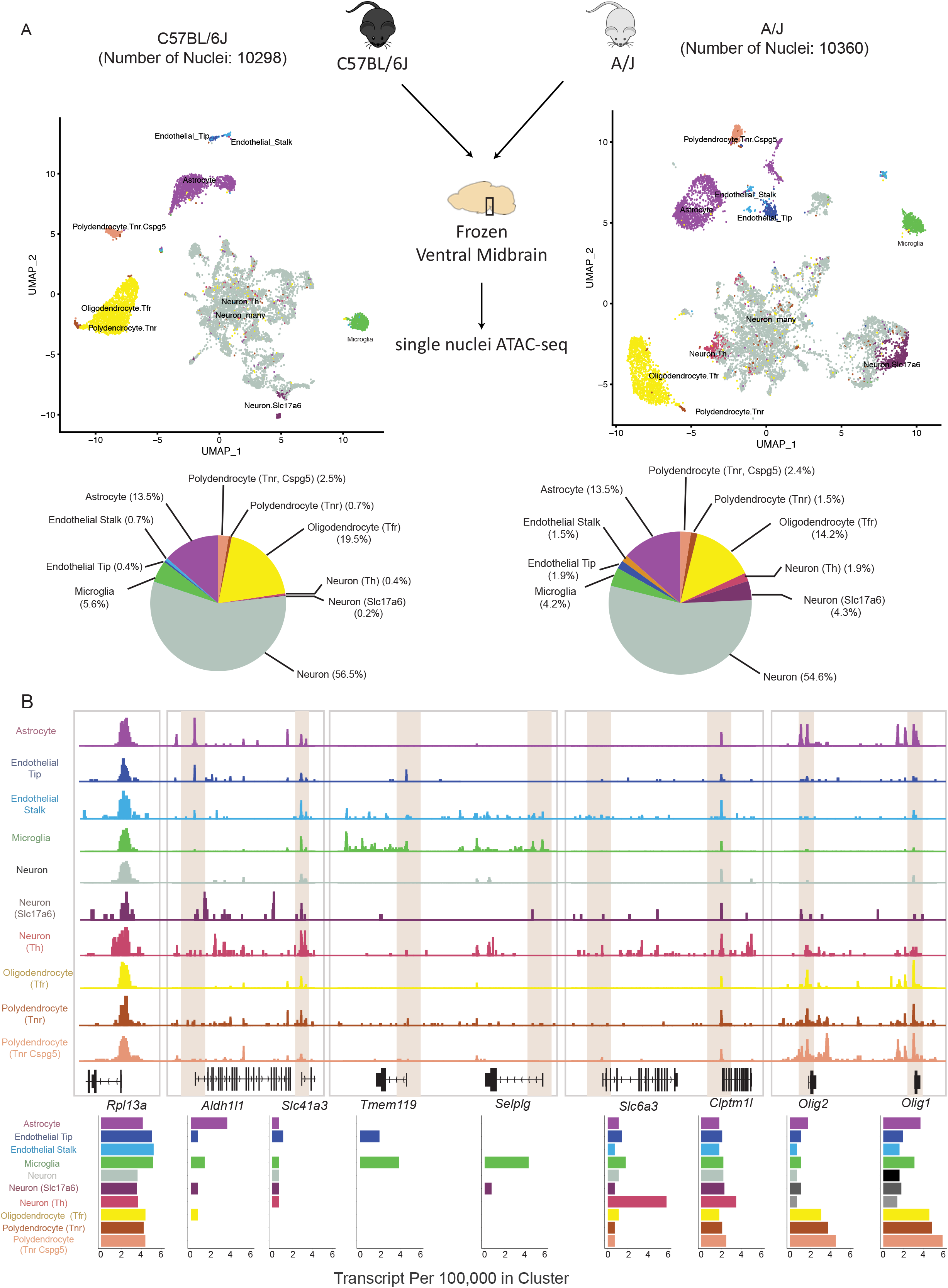
Midbrain snATAC-seq identifies cell type-specific accessibility. (A) Clustering of snATAC-seq from C57BL/6J and A/J with corresponding cell type proportions. Major cell types can be identified based on snATAC-seq profiles, with neurons having the biggest proportion on both strains. Cell types in C57BL/6J and A/J have comparable proportions with more than half of neuclei being identified as neurons. (B) Cell type-specific accessibility is observed in marker genes. The genomic tracks are from C57BL/6J midbrain snATAC-seq. The expression profiles measured as transcript per 100 000 in cluster. *Rpl13a* is used as a house keeping gene to normalize the snATAC-seq signal. See also Figure S1 and S2.

Using an existing single cell genomics toolkit (32, 33), the dimensionality of snATAC-seq was calculated by performing latent semantic indexing (LSI), to allow clustering of the cells with uniform manifold approximation and projection (UMAP) (Figure 1A). A gene activity matrix of snATAC-seq was established by counting reads in the gene body and the promoter region (2 kb upstream of transcription start site (TSS)). To annotate the obtained clusters as individual cell types, we took advantage of existing single cell RNA sequencing (scRNA-seq) of mouse midbrain (8). Through identification of anchor genes shared between the gene activity matrix of snATAC-seq and the highly variable features in scRNA-seq, we could identify 10 different midbrain cell types with distinct chromatin accessibility profiles and sufficient numbers of cells in both strains (Figure 1A, Table S1). The cell types are grouped into 6 main clusters consisting of glial cells like astrocytes (13.5%), microglia (4.2-5.6%), oligodendrocytes (14.-19.5%), and two subtypes of polydendrocytes (*Tnr^+^* and *Tnr^+^/Cspg5^+^*) (3.2-3.9%), two different types of endothelial cells (stalk and tip; 1.1-3.4%), and the largest and most diffuse cluster neurons making up more than half of all cells (57.1-60.8%). Although scRNA-seq data could distinguish up to 30 neuronal subtypes in midbrain through combinations of marker gene abundances (8), at chromatin accessibility level these could not be clearly distinguished. Instead, only three classes of neuronal cell types could be well distinguished: thalamus glutamatergic neurons (referred to as *Slc17a6^+^* neurons), dopaminergic neurons (referred to as *Th^+^* neurons), and a broader group of neurons consisting to large extent, but not exclusively, from different *Gad2^+^* GABAergic neurons (referred to simply as neurons).

Interestingly, an increased proportion of *Th^+^* and *Slc17a6^+^* neurons and decreased proportions of oligodendrocytes and macrophages could be detected in A/J samples compared to C57BL/6J, while the proportion of astrocytes and *Tnr^+^/Cspg5^+^* polydendrocytes remained almost identical (Figure 1A).

Inspection of genomic loci encoding for known cell type-specific marker genes in C57BL/6J samples disclosed highly cell type-selective chromatin accessibility that was well correlated with gene expression levels in scRNA-seq data of mouse midbrain (Figure 1B). While ubiquitously expressed *Rpl13a* gene had high and consistent levels of accessibility across the cell types, known marker genes for astrocytes (*Aldh1l1*) and microglia (*Tmem119* and *Selplg*) (34, 35) were expressed and most accessible in the respective cell types, especially at their TSS. Similarly, the gene encoding for dopamine transporter (*Slc6a3*) had highest levels of exression and accessibility in the *Th+* neurons, while in other neurons almost no signal could be detected. At the same time, the adjacent *Clptm1l* gene harboured an accessible promoter in all of the cell types. Finally, the locus encoding for two TFs required for oligodendrocyte generation and maturation, *Olig1* and *Olig2* (36, 37), showed highest accessibility in subtypes of polydendrocytes and oligodendrocytes, as well as astrocytes, again consistent with the gene expression levels.

Importantly, the accessibility profiles between C57BL/6J and A/J were highly comparable also at the level of individual cell types and could equally highlight cell type-specific accessibility consistent with gene expression levels, as shown in Figure S1 for *Aif1*, a known marker gene for microglia.

Taken together, our snATAC-seq profiling produced over 20 000 chromatin profiles of mouse midbrain cell types with comparable quality from two different mouse strains. These data allow the distinction of 10 different midbrain cell types at epigenomic level that are consistent with known gene expression profiles.

### Identification of cell identity genes and associated regulatory regions from single cell data

To leverage the available data for the identification of TFs controlling cellular identity in adult midbrain cell types, we first set out to determine the genes whose expression was selective for each cell type. To obtain these cell identity genes, we used the existing scRNA-seq of the mouse midbrain, and for each gene determined the 85^th^ percentile of its expression across all cell types indicating gene specific “high expression” (Figure 2). To filter out genes being selectively expressed in a specific cell type, at least 60% of the cells of that cell type have expression not less than the 85^th^ percentile. Furthermore, to ensure uniqueness, no other cell type was permitted to have the same gene among its top expressed genes in more than 40% of the cells. Through this approach we could define between 47 and 412 identity genes for each cell type (Figure 2; Table S2). On average, 170 genes per cell type were determined. To confirm the relevance of the genes to the biology of the cell type in question, GO enrichment analysis for biological processes was performed. In keeping with the genes’ role in the molecular and biological identity of the cell types, the top enriched GO terms included: Positive regulation of angiogenesis for endothelial stalk cells; Neutrophil mediated immunity for microglia; Axonogenesis, Neurotransmitter transport, and Regulation of synaptic vesicle exocytosis for the different neurons; Septin ring assembly and Myelination for oligo- and polydendrocytes; and Negative regulation of neuron differentiation for astrocytes. Examples of gene expression profiles of identity genes from selected cell types are shown in Figure 2B. Full list of enriched GO terms are provided in Table S3.

**Figure 2.**
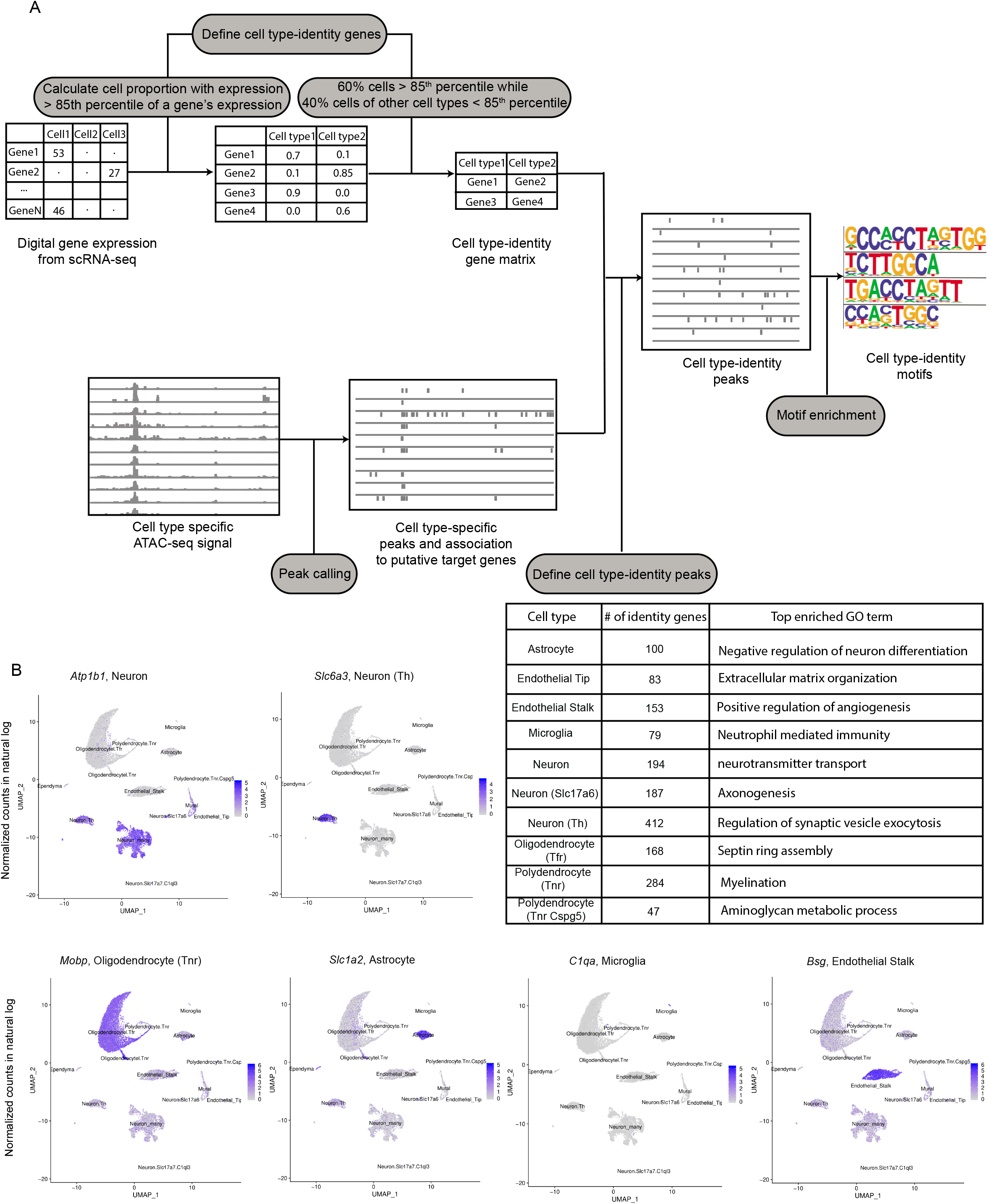
Regions controlling cell type identity can be defined by combining snATAC-seq and scRNA-seq. (A) Schematic workflow to define cell-type specific signatures. Digital gene expression is obtained from DropViz. For each gene, the 85^th^ percentile of its expression across all cell types was calculated. To define a gene as a cell type-identity gene, at least 60% of the cells of a cell type should have expression more than the 85^th^ percentile, while at the same time no other cell type was permitted to have the same gene among its top expressed genes (above the 85^th^ percentile) in more than 40% of the cells. Enrichment analysis with cell type-identity genes found GO terms corresponding to cell type characteristics. The cell type-identity peaks are defined by peaks overlapping with the regulatory regions of cell type-identity genes (basal region +/− 100 kb until nearby genes). Subsequently, the enriched motifs in cell type-identity peaks are detected. (B) Examples of identified cell type identity genes. The identified cell type-identity genes for six major cell types show selective expression in the respective cell types when observing scRNA-seq data of the entire population of midbrain cells.

Next, to determine gene regulatory regions controlling the expression of cell identity genes, we performed peak calling on cell type-specific aggregate ATAC-seq signals and associated the peaks to the defined identity genes of the respective cell types using GREAT (basal regulatory region +/−100 kb from TSS or up to nearest gene (38)). This resulted in 100 – 1200 accessible regions likely to control cell identity gene expression in each cell type (Figure 2; Table S4).

### Cell type-specific chromatin accessibility profiles uncover cell identity regulating transcription factors

Comparison of the chromatin accessibility levels across the cell types confirmed a clear increase in accessibility at the obtained cell identity peaks associated with respective cell identity genes (Figure 3A). The highest increase in signal over background of aggregated midbrain cells was always detected in the corresponding cell types expressing the associated identity genes. At the same time, depletion of signal could be detected in other cell types. Interestingly, the level of accessibility also reflected the developmental relationships of the cell types. The strongest depletion of signal could be detected in the developmentally most distant cell type, microglia (39). Consistently, microglia identity peaks showed the strongest depletion in all other cell types. In contrast, neuron identity peaks showed no major depletion of signal in the related *Th^+^* neurons and *vice versa*. Altogether, our approach could accurately identify cell type-specific gene regulatory regions controlling cell type identity.

**Figure 3.**
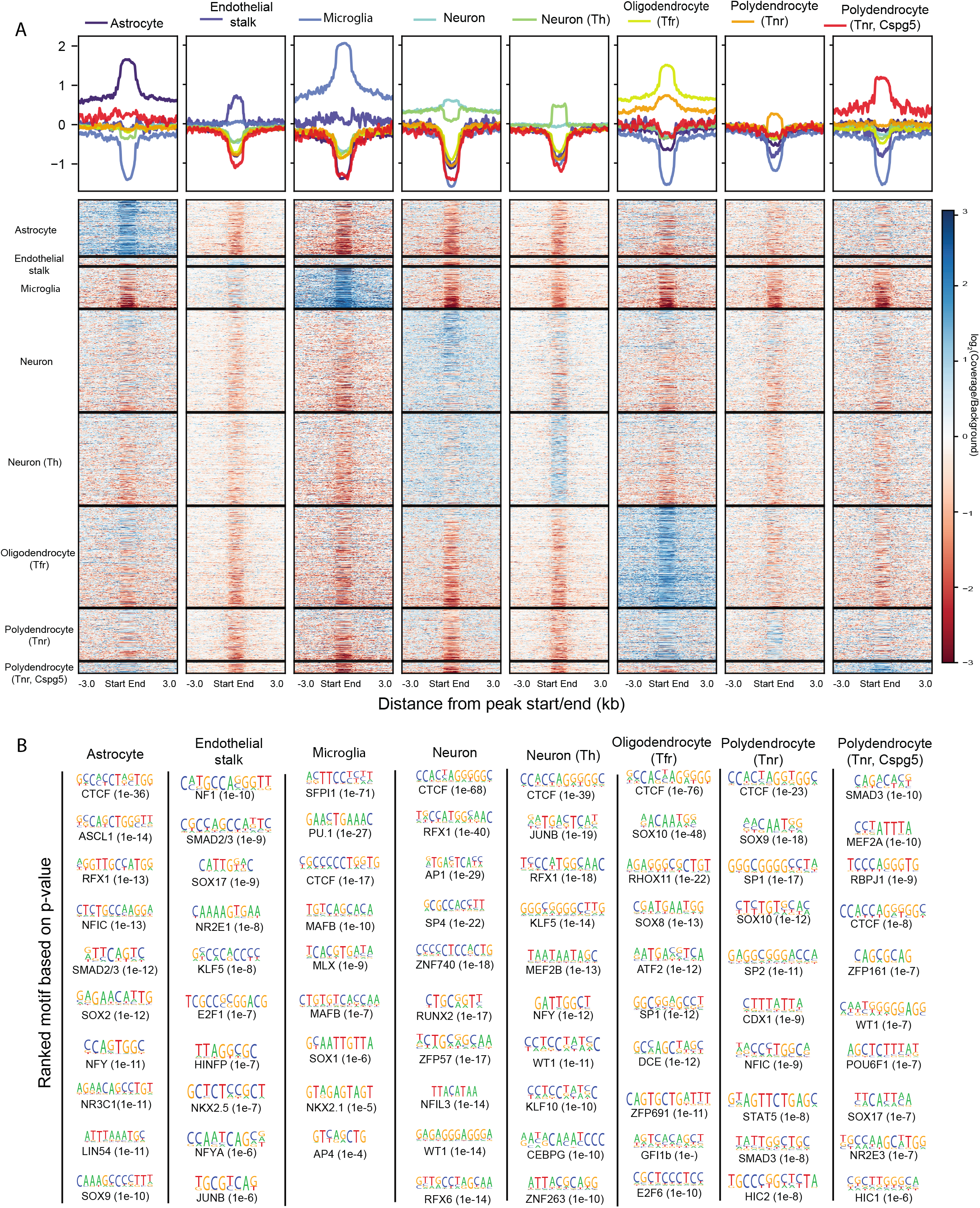
Identification of cell type specific TFs controlling cellular identity. (A) Heatmap showing the enriched signal of cell type-identity peaks in eight cell types. The analysis was done on C57BL/6J midbrain snATAC-seq. The background is constructed by merging the sampling reads (366278 reads / cell type) from each cell type. The raw signal is normalized to the background and library, following log_2_-transformation. The normalized signal is plotted 3 kb up- and downstream of peaks. (B) Motif enrichment analysis on cell type-identity peaks. The PWM logos, names of the associated TFs and p-values are shown for each motif. The motifs are ranked according to p-values.

To identify TFs binding the regulatory regions and controlling cell type identity genes, we performed TF binding motif analysis in sequences enriched at cell identity peaks (Figure 3B). This analysis was done for eight cell types with the highest sequencing coverage. Importantly, the analysis highlighted motifs for several TFs previously shown to control the differentiation or identity of the respective cell types. These included SOX9 in astrocytes (40), SPI1 in microglia (also known as SFPI1 or PU.1 (41, 42)), SOX17 in endothelial stalk (43), and SOX10 and SOX8 in oligodendrocytes (40, 44). The most enriched motif across cell types was the shared binding site for CTCF and CTCFL, the sequence occuring at insulator regions where CTCF mediates chromatin looping events (9), indicating the formation of cell type-selective chromatin looping and topological domains at the loci of cell identity gene loci.

In addition, NFI-family motif was enriched in astrocytes and polydendrocytes, consistent with its reported role in the transition from neurogenesis to gliogenesis (45) and the requirement of NFIC for expression of astrocyte marker genes (46). Motifs enriched in microglia included MAF motif which can be bound by MAFB, a TF recently shown to be important for maintenance of homeostasis in adult microglia (47). Interestingly, RFX-family motif was highly enriched in both astrocytes and neurons. Indeed, *Rfx1*, *Rfx3*, *Rfx4*, and *Rfx7* are known to be expressed and to play a role in the brain (48), with *Rfx4* showing the strongest expression in astrocytes while *Rfx3* and *Rfx7* are abundant in different neurons (8). Thus, our data warrant further investigation of individual TFs in the RFX-family in the cellular identity of midbrain neurons and astrocytes.

Together with RFX family, another TF with enriched motif in neurons, ZNF740, also has been shown to localize at gene enhancers active specifically in differentiated human neuronal cell lines, further supporting the relevance of this prediction across species (49). Finally, the motifs enriched uniquely in *Th^+^* neurons included binding sites for KLF family TFs, MEF2 TFs, and ZBTB7 TFs (that share their core motif with WT1 and ZNF263) (Figure 3B). From these particularly *Klf9, Mef2a, Mef2d*, and *Zbtb7c* show high expression in *Th^+^* neurons (8), with *Mef2d* exhibiting the most selective expression, an observation that could guide more detailed experiments into their role in dopaminergic neuron identity.

### Genetically driven chromatin accessibility changes reveal cell type-specific gene expression changes

We have recently shown that the midbrain phenotypic differences between C57BL/6J and A/J (and associated behavioural changes) are accompanied by extensive gene regulatory variation (20). Based on tissue-level bulk RNA-seq analysis, 1151 genes are significantly differentially expressed (>2-fold, FDR<0.05) in the ventral midbrains between the two strains (Figure 4A). However, the *cis*- and *trans*-acting mechanisms underlying these genetically driven changes, and the affected cell types, are not known.

**Figure 4.**
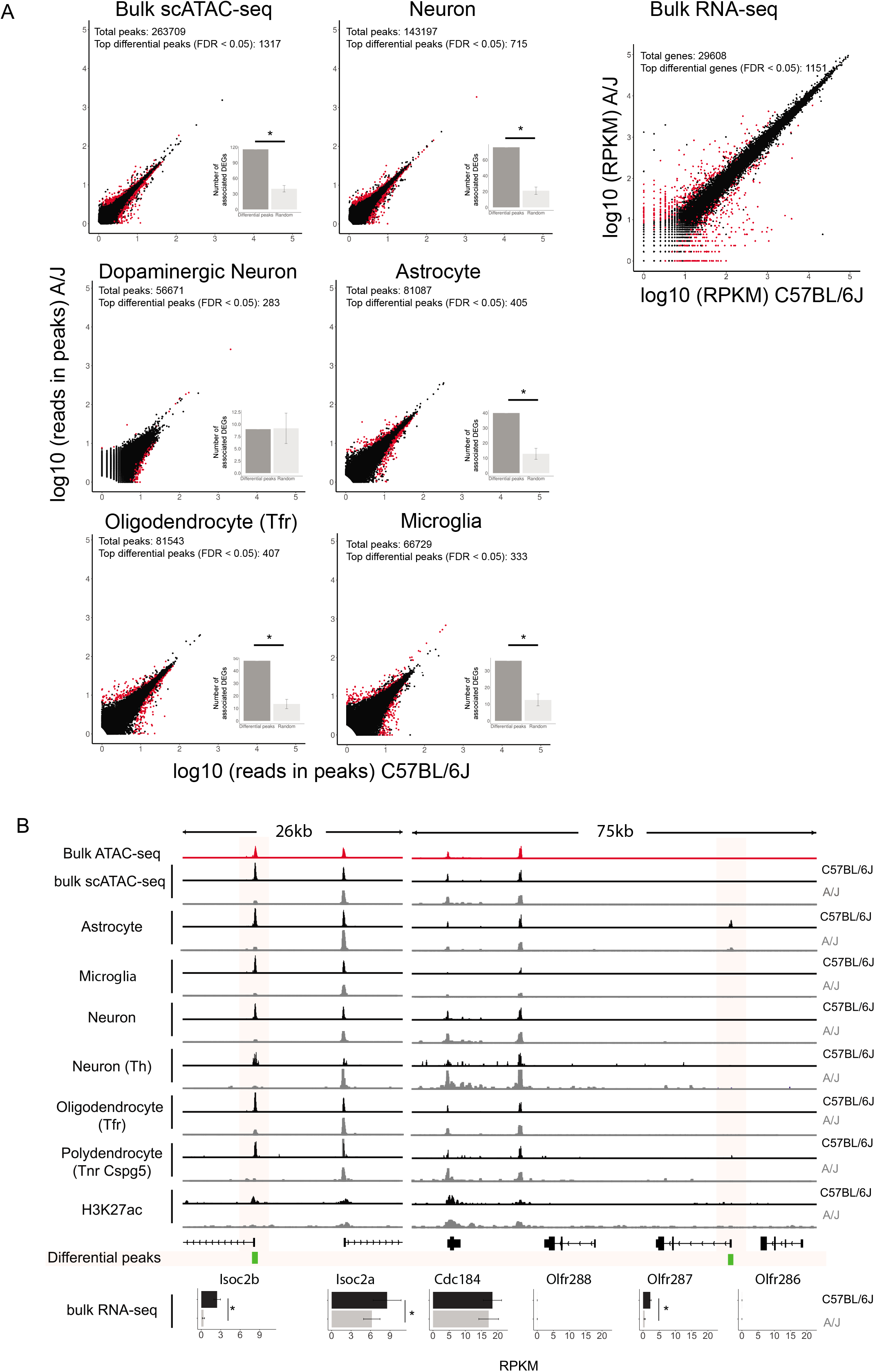
Association of differentially accessible regions with altered gene expression between C57BL/6J and A/J. (A) Differential peaks are highly associated with differential genes. Top differential peaks (labelled as red) are defined by Wilcoxon rank-sum test with FDR < 0.05 within top 1% of logFC. The read counts in peaks of snATAC-seq bulk are log_10_-transformed. Peaks with low read count (less than median - 1.5 median absolute deviation) are filtered out. To associate differential peaks to DEGs, peaks are overlapped with the regulatory region of DEGs (basal region +/− 100 kb until nearby genes). As a control, random peaks are selected by bootstrapping with 1000 repetitions (p < 0.0099). The RPKM from bulk RNA-seq of C57BL/6J and A/J is also log_10_-transformed, and DEGs are defined as FDR < 0.05 and log_2_-fold change > 1 (labelled as red). (B) Cell type-specific differential peaks correlate with gene expression in bulk RNA-seq. The differential peaks are labelled as green.

To address the contribution of chromatin level changes to the gene expression variation, we compared the bulk snATAC-seq signals from the mouse strains and focused on top differential peaks (FDR < 0.05, top 1% of logFC) (Figure 4A). The extent of fold changes at the chromatin accessibility level was more modest than the transcriptomic level. Nevertheless, 1317 of 263 709 called peaks were significantly altered at the bulk level. By associating accessible regions to their target genes, we could observe a significant enrichment of such regions at the differentially expressed genes. This indicates at least some of the gene expression changes could be linked to changes at the chromatin level. Observing the data at the level of individual cell type allowed detection of additional cell type-specific differentially accessible regions, indicating that cell type-specific changes from rare cells could be masked by tissue level analysis. The significant enrichment of the differential peaks at differentially expressed genes held in all cell types except dopaminergic neurons which might result from their low sequencing coverage (Figure 4A).

For genes like *Isoc2b*, the decreased gene expression in ventral midbrain of A/J was accompanied by reduced accessibility of the promoter across all cell types (Figure 4B). To confirm that the lost accessibility was also accompanied by reduced transcriptional activity, we observed H3K27ac levels at the promoter. Consistent with reduced ATAC-seq signal, H3K27ac was also lost at *Isoc2b* locus in A/J.

For ubiquitously expressed genes like *Isoc2b* the altered gene expression could be associated with chromatin level changes even at bulk level analysis. However, for other genes such as *Olfr287*, a reduced expression could be observed by RNA-seq although no signal was detectable at bulk chromatin level by any of the methods (bulk ATAC-seq, bulk snATAC-seq, and ChIP-seq). Still, when observing the cell type-specfic snATAC-seq data, an accessible region could be detected at *Olfr287* promoter specifically in astrocytes. And consistently with reduced gene expression, the chromatin was less accessible in A/J (Figure 4B).

In summary, gene regulatory variation in midbrain is associated with chromatin level changes in accessibility, although not at all loci and with lower sensitivity than in transcriptomic analysis. Interestingly, snATAC-seq can reveal cell type-specific regulatory changes not captured in bulk level analysis.

### Putative *cis*-acting variants are enriched at midbrain regulatory regions of differentially expressed genes

To obtain further insight into the mechanisms underlying the strain-specific gene expression, we next set out to address the extent of *cis*-acting regulatory variation contributing to the observed differences in the midbrain. For this we focused on identification of putative midbrain regulatory variants segregating C57BL/6J and A/J. We first performed TF footprint identification on our midbrain chromatin accessibility profile obtained through the bulk ATAC-seq analysis. Then, these footprints were overlapped with >6 million variants segregating C57BL/6J and A/J to identify those with the potential to disrupt TF binding. Finally, the binding sites were overlapped with the midbrain H3K27ac profiles from both C57BL/6J and A/J to capture the binding sites engaged in transcriptional activity in either mouse strain, in total yielding 3909 putative regulatory variants of the ventral midbrain (Table S5).

The capacity of the above approach to reduce the number of meaningful variants is illustrated in Figure 5A with the examples of the *Ddhd1, Zfp615*, and *4.5S rRNA* loci. Expression of *Ddhd1*, a gene coding for a phospholipase, is modestly but significantly reduced in A/J compared to C57BL/6J and shows accessible chromatin at its TSS and at an upstream enhancer site >20 kb from the TSS. Both regions are marked by H3K27ac signals in both strains. One TF footprint could be identified at both the TSS and the distal enhancer, representing the putative TF binding sites controlling *Ddhd1*. From total of 603 variants at the 61 kb locus, only one coincides with an active TF binding site occupied in the midbrain, corresponding to a putative regulatory variant influencing *Ddhd1* expression in this brain region. Consistently, the affected enhancer shows decreased H3K27ac enrichment in the A/J. This illustrates how majority of genetic variants at any given locus are unlikely to affect gene expression and how, by focusing on those co-localizing within active regulatory regions, those most likely to act as regulatory variants can be identified.

**Figure 5.**
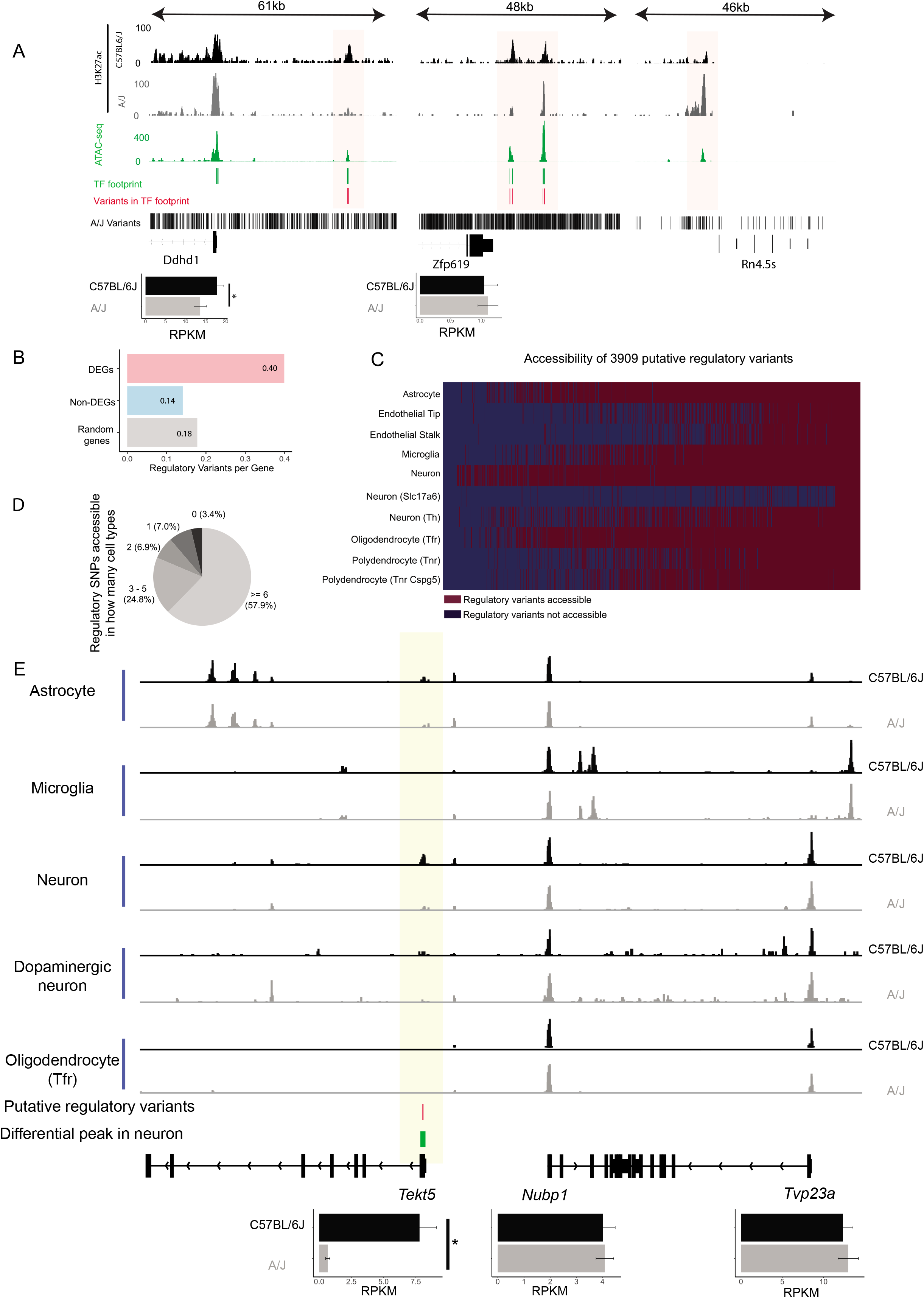
Putative regulatory variants are associated with differentially expressed genes and show cell type-selective accessibility. (A) Examples of putative regulatory variants of A/J found in the enhancer region upstream of *Ddhd1, Afp619* and *Rn4.5s*. The putative regulatory variants are defined as variants disrupting TF footprints located in active enhancers (defined by H3K27ac). (B) Each DEG (5082, FDR > 0.05) is associated with an average of 0.4 putative regulatory variants, while non-DEGs are associated with only 0.14 variants. Random: 5000 genes are randomly selected from all expressed genes. (C) Putative regulatory variants have differentical accessibility across cell types. Differential accessibility of putative regulatory variants is shown in the heapmap (red: accessible, blue: non-accessible). (D) More than half of the variants are accessible in more than 6 cell types while 7% in only 1 cell type. (E) An example showing how putative regulatory variants with differential accessibility affect cell type-specific gene expression. The variants locating near the TSS of *Tekt5* are associated with TSS signal in neurons of C57BL/6J but not A/J, potentially resulting in upregulation of *Tekt5* as shown in bulk RNA-seq.

If the midbrain gene regulatory differences between C57BL/6J and A/J indeed depend on the cumulative effect of *cis*-acting variants, such as the variant at the *Ddhd1* locus, the identified regulatory variants would be expected to be enriched in regulatory regions and TF binding sites at the differentially expressed gene loci compared to other expressed genes. To test this directly, we associated all putative regulatory variants to their likely target genes as already outlined in Figure 2, and calculated the number of variants that on average associate with each of the 5082 differentially expressed genes (FDR<0.05) (Table S6) (20). We did the same for all expressed genes found not to change between the strains (FDR>0.05) and for an equal number of randomly selected expressed genes as controls. While unaffected genes and randomly selected genes were associated on average with 0.14 and 0.18 regulatory variants, respectively, this number significantly increased to 0.40 regulatory variants for the differentially expressed genes (Figure 5B). Consequently, genetic variants located in midbrain regulatory regions do not show a random distribution but are instead enriched at the differentially expressed genes, suggesting they play an important role in explaining the observed transcriptomic differences.

Next, we considered whether localization of variants in the TF binding sites of active enhancers could also be associated directly with enhancer activity upstream of gene expression changes. With this aim we used THOR (29) to identify enhancer regions with significantly altered signal for the H3K27ac enhancer mark between midbrains of C57BL/6J and A/J. Interestingly, 1126 of the 3909 putative regulatory variants localized within an enhancer region with differential H3K27ac enrichment under a stringent cut-off (p<1×10^−18^) (Table S7). This indicates that a large proportion of putative regulatory variants associate with enhancers that gain or lose activity between the mouse strains. For example, enhancer harbouring putative regulatory variants in the proximity of *4.5S rRNA* locus exhibits a strong gain of enhancer activity in A/J compared to C57BL/6J (Figure 5A). And at locus like *Zfp619* both gain and loss of enhancer activity can be observed simultaneously at two separate enhancers associated with multiple putative regulatory variants. Taken together, disruption of TF binding by variants across thousands of enhancer regions is likely to alter enhancer activity, and thereby midbrain gene expression in genetically diverse mouse strains.

### Cell type-specificity of *cis*-acting variants in the midbrain

Having identified the putative *cis*-acting regulatory variants contributing to the midbrain gene expression phenotype between C57BL/6J and A/J, we next sought to understand how cell type-selective these variants are. Overlapping the putative regulatory variants with cell type-specific accessibility data suggested that majority of the variant binding sites (57.9%) were accessible, with potential to affect gene expression, in at least 6 out of the 10 cell types (Figure 5C-D). However, just under 14% of the variants were accessible in only 1 or 2 cell types, indicating non-coding variation can have cell type-selective effects on gene expression (Figure 5D). Indeed, variants with cell type-selective accessibility in only 1-3 cell types were significantly more often occurring at genes with altered expression than at other expressed genes (data not shown).

A number of regulatory variants were accessible specifically in neurons and associated with differentially expressed genes, representing neuron-specific gene regulatory variation. These include for example *Tekt5*, a gene expressed in excitory neurons and upregulated in C57BL/6J (Figure 5E). Despite the observed differential expression, *Tekt5* promoter appeared inaccessible in most cell types except for neurons of C57BL/6J, exactly at TSS overlapping with a putative regulatory variant within a TF binding site.

Taken together, while majority of *cis*-acting variants affect broad array of cell types, a large proportion can have cell type-specific effects that cannot be dissected without single cell analysis.

### TCF7L2 as a mediator of *trans*-acting variation in midbrain neurons

A large fraction of midbrain gene expression variation could be linked to *cis*-acting regulatory variants, even with our strict criteria on the presence of variant in a TF footprint located in an active enhancer (Figure 5A). Still, much differential gene expression remained unexplained. This could be due to *cis*-acting variants we have missed, but also due to *trans*-acting variants that can influence a number of target genes by altering a TF’s activity, rather than its binding motif. A change in TF activity could be due to change in its expression level, but could also be due to alternations in other mechanisms controlling TF activity such as post-translational modifations, protein-protein interactions or TF localization.

Genetic differences in non-dopaminergic neurons (such as *Gad2^+^* neurons), that make up much of our neuron population (Table S1), have been suggested to contribute to strain specific behavioural differences, including anxiety, reward and motivation traits, such as ethanol consumption (50–53). To identify mediators of *trans*-acting variation between C57BL/6J and A/J in neurons, we performed motif enrichment analysis on 715 regions in neurons showing differential chromatin accessibility between the mice (Figure 4A). This revealed the binding motif of TCF/LEF family, downstream TFs of the canonical Wnt signaling pathway (54), to be among the most enriched sequences found at the differentially accessible regions (Figure 6A; p = 1.66e-14). The enrichment was specific for *Gad2^+^* neurons and could not be found in any other cell types (Figure S3) or in the motif enrichment analysis for cell identity genes (Figure 3). LEF1, TCF7L1, and TCF7L2 bind to the same DNA sequence but have often opposing or cell type-specific functions (55). Inspecting chromatin accessibility across the cell types for the differential binding sites carrying the TCF/LEF motif revealed an increased signal specifically in the neurons (Figure 6B). To determine which factor is expressed in the midbrain neurons and could mediate the observed enrichment and altered accessibility, we visualized their expression using scRNA-seq data. *Lef1* expression was limited to the endothelial cells (Figure 6C) and *Tcf7l1* showed only low or no expression across the cell types (Figure 6D). However, *Tcf7l2* had high expression in the *Gad2^+^* and *Slc17a6^+^* neurons and polydendrocytes, showing a clear overlap with cells enriched for the respective binding motif (Figure 6D). Consistently, an analysis of putative upstream regulators explaining the transcriptomic changes from RNA-seq of the mouse strains using Ingenuity Pathway Analysis (IPA) predicted TCF7L2 (also know as TCF4) and CTNNB1 (beta-catenin, binding partner of TCF7L2) to be among the top regulators based on the predicted activation score (Table S8). Thus, the transcriptional activity of TCF7L2 is likely to be altered between C57BL/6J and A/J mice in the midbrain neurons.

**Figure 6.**
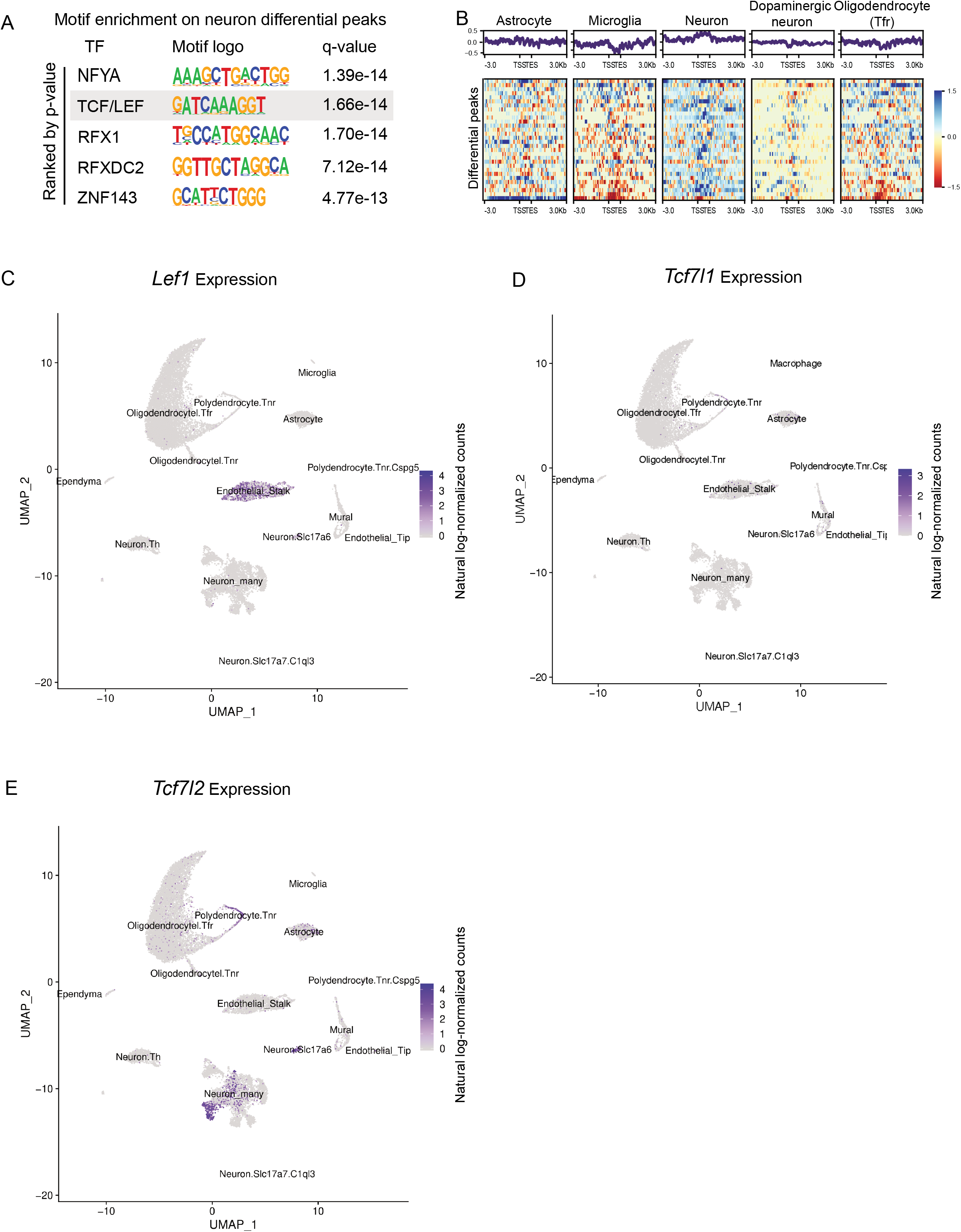
TCF/LEF family as a mediator for *trans*-acting variation in neurons. (A) Motif enrichment analysis for regions of differential accessibility between C57BL/6J and A/J in neurons. (B-C) Regions of differential accessibility with motif for TCF/LEF family show increased accessibility in neurons compared to other cell types. (D-E) *Lef1* is highly expressed in endothelial stalk*, Tcf7l1* shows low expression in all cell types, while *Tcf7l2* is abundant in neurons and polydendrocytes. See also Figure S3.

Taken together, snATAC-seq analysis of tissues from genetically different strains can guide the elucidation of cell type-specific impact of *trans*-acting variants and suggests neuron-specific differences in the canonical Wnt signalling pathway between two commonly used inbred mouse strains.

## DISCUSSION

Here we investigated the chromatin accessibility of cell types in mouse ventral midbrain in two different genetic backgrounds and provided a large resource of 20 658 single nuclei chromatin profiles on 10 different cell types. This dataset will benefit future studies on the role of these cell types in various processes involving ventral midbrain such as movement control, cognition and reward mechanisms. A better understanding of midbrain cell types can also profit research on diseases like PD and schizophrenia. In particular, improved comprehension of genetic variation in gene regulation and how it impacts specific cell types, will pave the way for better prediction of genetic susceptibility and affected disease mechanisms.

Our findings on gene accessibility profiles and cell type composition of the midbrain are consistent with existing knowledge from single cell transcriptomics (8). However, neuronal subtypes are more difficult to distinguish at chromatin level than what has been achieved by transcriptomic analysis. Neuron subtypes clustered largely together, often with undefined borders between the subtypes (Figure 1A). This result is expected. While chromatin accessibility is generally known to show positive correlation with gene expression (56, 57), and this is also true for our data (Figure S2), enhancer accessibility does not necessarily reflect gene regulatory activity (58). Indeed, chromatin accessibility profiles of cell types executing similar functions can be highly similar despite showing different expression patterns and being controlled by different master TFs (16, 59). Moreover, accessibility can signify priming of a locus for expression without commencing transcription (60, 61).

Nevertheless, we could reliably distinguish 10 of the 20 known midbrain cell types at the chromatin level, with 8 cell types containing sufficient cells for detailed analysis (Figure 3). Using the information about gene regulatory regions selectively associated with genes underlying cellular identity, we were able to predict the key regulators of each cell type through motif enrichment analysis. These included many factors previously determined to be necessary for the differentiation or the maintenance of the respective cell state (40–42, 44, 62). Among other insights, the results reveal a cell type-specific role for CTCF binding sites in cellular identity and predict specific roles for MEF2, KLF, and ZBTB7 family TFs in dopaminergic neurons. Moreover, the results give additional support for a more detailed analysis of RFX family of TFs in regulation of neuro- and gliogenesis balance in the midbrain, something that is consistent with previously identified co-occupancy of RFX factors at SOX2-bound enhancers in neurodevelopment (63).

Genetic variation is known to drive gene expression changes that can trigger phenotypic differences (64–67). We have recently shown over thousand genes to be differentially expressed in the mouse midbrain due to genetic variation between C57BL/6J and A/J (20). Here we leveraged our single cell chromatin accessibility profiles to investigate the mechanisms, cell type-specificity, and extent to which this variation is reflected at the level of chromatin. Interestingly, the transcriptomic differences did not show a linear relationship with midbrain chromatin accessibility (Figure 4A). The extent of change in accessibility was not comparable to mRNA level change either at bulk aggregate levels or in individual cell types. This is most likely reflecting the observation that TF binding and activity can alter gene expression without change in accessibility, for example if the locus is already open (68). Still, when changes in accessibility did occur, this was often associated with local gene expression change. Importantly, some of the chromatin accessibility changes associated with differential gene expression could not be detected at all in tissue level ATAC-seq or ChIP-seq analysis despite high sequencing depth (Figure 4B). Therefore, the improved resolution offered by single cell analysis allows further insights into regulatory interactions that could be missed in tissue level analysis.

*Cis*-acting variants that influence gene expression have been suggested to explain a significant part of the missing heritability (69–71). Increasing number of such variants associated with human traits and diseases have now been experimentally validated (17). By combining genetic information with ATAC-seq and ChIP-seq analysis, we found 3909 of the >6 million variants segregating C57BL/6J and A/J to localize in a TF binding site within an active and accessible enhancer in the ventral midbrain (Figure 5). This number is consistent with the previous estimates that one in thousand mouse variants cause *cis*-regulatory effects (72). Importantly, we found the putative *cis*-acting variants to be enriched at differentially expressed genes, indicating they do contribute to the gene expression phenotype.

A large proportion of the putative *cis*-acting variants showed cell type-selective accessibility (Figure 5B). This finding is consistent with previous work on impact of *cis*-acting variants on disease-associated regulatory variation in the major cell populations of human brain (73). Combining the information on cell type-specific impact of variants on gene regulation with genome-wide association studies and quantitative trait locus mapping for specific traits or diseases can provide insights on how the variants translate into phenotypes. C57BL/6J and A/J differ from each other for numerous phenotypes (74). These include fear-conditioning and reward-related behaviours like ethanol consumption that are associated with midbrain dopamine signalling (51–53). Combining our data on cell type selectivity of *cis*-acting variants with genome-wide association studies of such phenotypes could reveal new connections between genes and the underlying mechanisms.

Besides *cis*-acting variation, differential gene expression and altered chromatin accessibility can also be mediated through *trans*-acting variation. *Trans*-acting variants can have many different mechanisms of action and the majority of genetically driven gene expression variation has been suggested to be due to *trans*-acting variation, often affecting hundreds or thousands of genes in a cell type-specific manner (70, 75, 76). Dopaminergic neurons in ventral tegmental area of the midbrain contribute to phenotypes like reward-behaviour and fear (50). Moreover, the associated GABAergic and glutamatergic neurons in midbrain can also control such behaviours, either together with or independent of dopamine signalling. With this in mind, we asked whether specific signature of *trans*-acting regulatory variation could be found in neurons between our strains that show differential behaviour, based on motif enrichment analysis at regions of differential accessibility (Figure 6). We found an enrichment for the TCF/LEF binding motif, corresponding to the Wnt signalling response element recognized by their high mobility group (HMG) DNA-binding domain (54). This suggests that Wnt signalling is altered between C57BL/6J and A/J in neurons. Based on scRNA-seq data, TCF7L2, but not LEF1 or TCF7L1, is abundantly expressed in midbrain *Gad2^+^* and *Slc17a6^+^* neurons, indicating that alterations in chromatin accessibility and gene expression are due to changes in Wnt signalling and likely to be mediated by TCF7L2 (Figure 6). Indeed, pathway analysis of the differentially expressed genes highlighted an enrichment of known Wnt target genes among the DEGs (Table S8).

Involvement of TCF7L2 as the putative effector of the altered signalling in neurons is particularly intriguing since *TCF7L2* locus has been genetically associated with mental disorders like scizophrenia in humans (77). Moreover, transgenic mice have revealed a dose-dependent role of *Tcf7l2* in fear-conditioning and anxiety, traits for which A/J is known to significantly differ from C57BL/6J (51, 52).

Understanding how Wnt signalling activity is altered between the mouse strains requires further analysis. However, it is interesting to note that expression of *Wnt2b*, upstream ligand of Wnt pathway genetically associated with bipolar disorder (77), is significantly decreased in the ventral midbrain of A/J compared to C57BL/6J (20).

Taken together, our single nuclei chromatin analysis provides novel insights into transcriptional control of ventral midbrain cell types and a rich resource for further analysis of cellular identity and gene regulatory variation in this disease-relevant brain region.

## DECLARATION

The authors declare no competing interests.

## DATA AND CODE AVAILABILITY

Raw FASTQ files were deposited in ArrayExpress with the accession number E-MTAB-8333 for bulk levels analysis and E-MTAB-9225 for single cell data. The snATAC-seq tracks are available at the UCSC Genome Browser (http://genome-euro.ucsc.edu/cgi-bin/hgTracks?hubUrl=https://biostat2.uni.lu/ygui/hub.txt&genome=mm10). The scripts for data analysis can be accessed here: https://github.com/sysbiolux/.

## SUPPLEMENTARY INFORMATION

Supplementary Information is available as separate Supplementary Files.

## ACKNOWLEDGEMENTS

We would like to thank Drs Aurélien Ginolhac and Anthoula Gaigneaux for their support with bioinformatic analysis, Dr Djalil Coowar (Animal Facility of University of Luxembourg) for help with breeding of experimental mice, Dr Rashi Halder at LCSB sequencing facility for high-throughput sequencing, and Sergio Helgueta for help with nuclei isolation. The computational analysis presented in this paper were carried out using the HPC facilities of the University of Luxembourg.

## AUTHOR CONTRIBUTIONS

YG, MB, and LS conceived the project. YG, KG, JO, AS, TS and LS designed the experiments. YG and KG performed snATAC-seq. YG and JO performed bulk ATAC-seq and YG performed ChIP-seq experiments. MT and PG prepared mouse tissues. YG performed all bioinformatic analysis with help from TS and LS. YG and LS analysed the results. MB, TS, and LS provided funding. YG and LS wrote the manuscript. All authors read and approved the final manuscript.

