## Supplemental Figures for "Single nuclei chromatin profiling of ventral midbrain reveals cell identity transcription factors and cell type-specific gene regulatory variation"

### Supplemental Information

### Supplemental Figures

**Figure S1: snATAC-seq on ventral midbrains of C57BL/6J and A/J revealed cell type-specific chromatin accessibility.**

A. The ventral midbrains of the two mouse strains were used as input to snATAC-seq. Ten cell types were identified based on clustering of peak features. The aggregated signal is comparable to bulk ATAC-seq. Different accessibility across cell types can be observed and they can reflect gene expression changes. *Prrc2a* is globally accessible and its expression can be detected in all cell types. *Aif1* and *Lst1* TSS are selectively accessible in macrophage, and their expression is only abundant in macrophage.

**Figure S2: H3K27ac ChIP-seq and ATAC-seq correlating with gene expression.**

A. H3K27ac ChIP-seq on ventral midbrains of C57BL/6J and A/J. Within-sample normalization is applied to account for gene length. The intensity of H3K27ac ChIP-seq signals are plotted in a window of 2000 bp upstream and downstream of gene body. The genes are ordered based on the highest to the lowest gene expression level.

B. ATAC-seq on ventral midbrain of C57BL/6J. The plotting scheme is the same as Supplementary Figure 2A.

**Figure S3: Differential peaks can reveal strain-specific TFs.**

A. Motif enrichment analysis on differential peaks. The PWM logos, names of the associated TFs and p-values are shown for each motif. The motifs are ranked according to p-values.

Figure S1

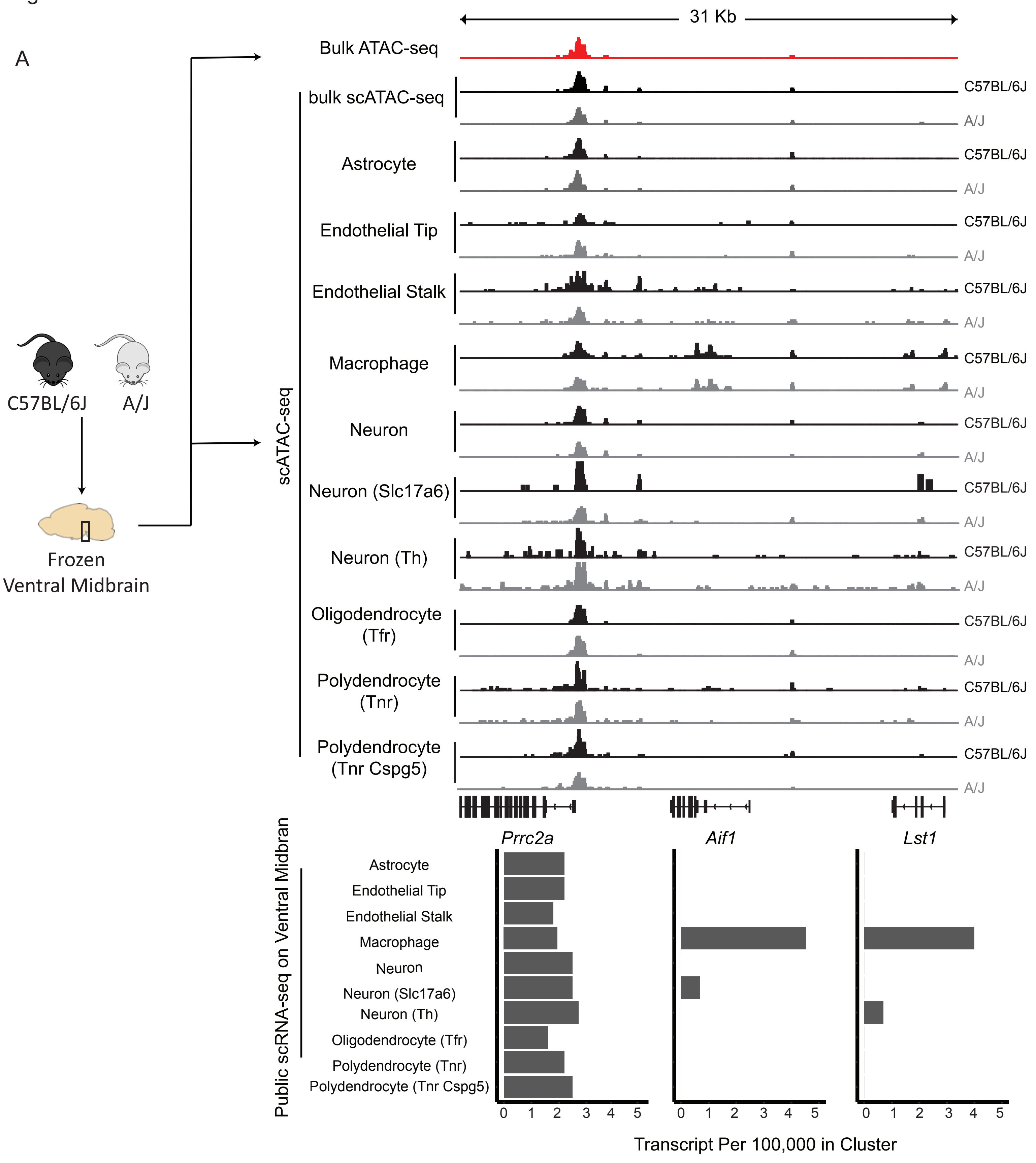
A
31 Kb
Bulk ATAC-seq
bulk scATAC-seq
Astrocyte
Endothelial Tip
Endothelial Stalk
Macrophage
Neuron
Neuron (Slc17a6)
Neuron (Th)
Oligodendrocyte (Tfr)
Polydendrocyte (Tnr)
Polydendrocyte (Tnr Cspg5)
C57BL/6J
A/J
scATAC-seq
C57BL/6J
A/J
Frozen Ventral Midbrain
Prrc2a
Aif1
Lst1
Public scRNA-seq on Ventral Midbran
Astrocyte
Endothelial Tip
Endothelial Stalk
Macrophage
Neuron
Neuron (Slc17a6)
Neuron (Th)
Oligodendrocyte (Tfr)
Polydendrocyte (Tnr)
Polydendrocyte (Tnr Cspg5)
0 1 2 3 4 5
Transcript Per 100,000 in Cluster

Figure S2

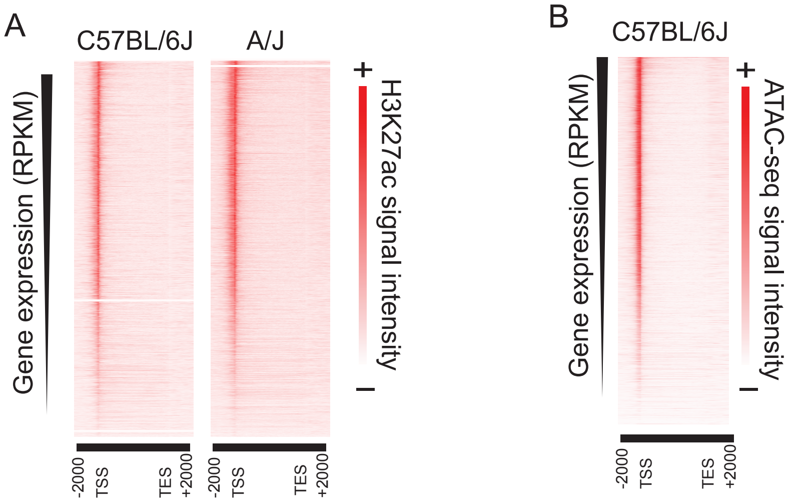

A
C57BL/6J
A/J
Gene expression (RPKM)
H3K27ac signal intensity
+
-
-2000
TSS
TES
+2000
-2000
TSS
TES
+2000
B
C57BL/6J
Gene expression (RPKM)
ATAC-seq signal intensity
+
-
-2000
TSS
TES
+2000

Supplementary figure 3

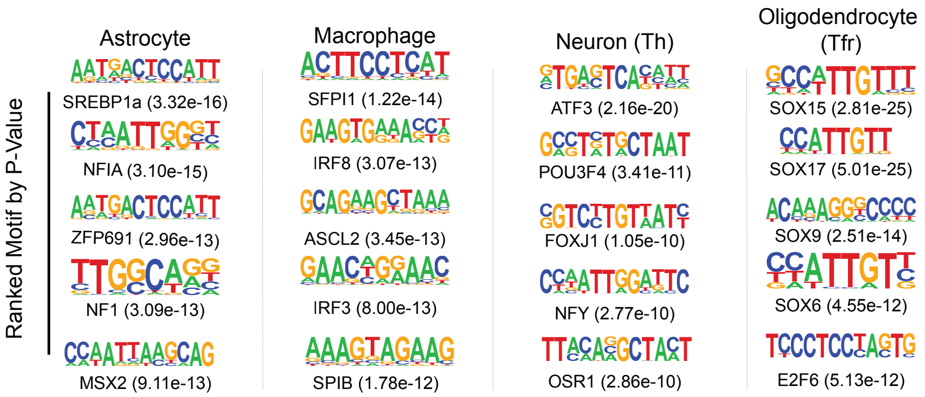

### Supplemental Tables

**Table S1: Cell type composition in ventral midbrains of C57BL/6J and A/J in snATAC-seq.** Cell types and identified cell numbers are indicated for both strains

**Table S2: Cell type-identity genes defined from existing scRNA-seq.** The official gene symbols for the identified cell identity genes are listed under each respective cell type.

**Table S3: Enrichment analysis on the cell type-identity genes defined from existing scRNA-seq.** The GO enrichment results for the cell identity genes for each cell type are provided. The enrichment analysis was performed using Enrichr and GO term identifiers, enrichment p-values, scores, and identified gene names are provided for each GO term.

**Table S4: Cell type-identity peaks defined by associating cell type-specific peaks to the regulatory regions of cell type-identity genes.** The chromosome coordinates for all cell identity peaks are provided for each cell type.

**Table S5: Putative regulatory variants.** The chromosome coordinates and major / alternative alleles are reported for the 3909 variants locating within TF footprints in active enhancers and designated as putative regulatory variants.

**Table S6: Differentially expressed genes from ventral midbrain bulk RNA-seq between C57BL/6J and A/J.** Each strain has 12 replicates (6 males and 6 females). The differential genes are defined as padj $< 0.05$.

**Table S7: H3K27ac differential peaks between C57BL/6J and A/J.** The chromosome coordinates of the peaks and p-values of signal difference between the strains are indicated for each differential peak ($p<1x10^{-18}$).



**Table S8: Predicted upstream regulators between C57BL/6J and A/J.** The predicted upstream regulators are fetched from IPA based on differentially expressed genes from midbrain bulk RNA-seq between C57BL/6J and A/J (padj < 0.05, log2FC > 1).
